# Early Prediction of Ischaemic Stroke Outcomes: A Novel Computational Approach

**DOI:** 10.1101/2024.09.25.615002

**Authors:** Xi Chen, Wahbi El-Bouri, Stephen Payne, Lei Lu

## Abstract

Malignant stroke can lead to a death rate as high as 80%. Although early interventions can improve patient outcomes, they also lead to side effects. Therefore, the early prediction of stroke outcomes is crucial for clinical strategies. Imaging markers such as brain swelling volume and midline shift have been critical predictors in various stroke scoring systems. However, these markers can only become visible on brain images days after stroke onset, which delays clinical decisions. A primary challenge in predicting these markers is that brain swelling is a biomechanical process that relies on anatomical features, such as lesion size and location. To tackle this problem, we propose a novel computational pipeline to predict brain swelling after stroke. We first provide a mathematical model of the brain by using a five-compartment poroelastic theory. It allows us to generate high-quality stroke cases with varied 3D brain and lesion anatomy, which are then used to train and validate a deep neural network (DNN). Our in silico experimentation with 3,000 cases shows that anatomical features of stroke brains are well-learned by the DNN, with minimal errors in brain swelling prediction found in the hold-out testing cases. In addition, we used the DNN to process clinical imaging data of 60 stroke patients. The results show that the markers generated from the DNN can predict 3-month stroke outcomes with an AUC of around 0.7. It indicates that the proposed computational pipeline can potentially advance the time point for clinical decisions.

**Significance Statement:** Stroke is the second leading cause of death in the world, and malignant stroke can lead to a death rate of 80%. Early interventions can improve patient outcomes but can also cause side effects. Therefore, it is crucial to predict stroke outcomes at an early stage. Radiological markers such as brain swelling volume and midline shift have been used in various stroke scoring systems. However, these markers can only become visible after days to stroke onset, which delays clinical decisions. To tackle this issue, we propose a novel computational pipeline to predict brain swelling after stroke onset. The proposed pipeline is found to predict brain swelling accurately and can potentially assist early clinical decision-making.

---

Stroke has been the second leading cause of death in the world (1), and one out of four adults aged over 25 will develop stroke in their lifetimes (2). Among the two types of strokes, ischaemic stroke is the most common (85%) (1), whereas haemorrhagic stroke consists of around only 15%. In ischaemic stroke, occlusion of the brain vessel leads to ischaemia and the degradation of the blood-brain barrier (BBB). As a result, the blood components leak into the interstitial space through damaged BBB and give rise to brain oedema. Malignant stroke, characterised by a large infarct territory and severe swelling of brain tissue, can lead to a mortality rate of 80% (3).

Early interventions have been effective ways to improve stroke outcomes (4). However, treatments such as osmotherapy (5) and hypothermia (6) can cause strong side effects that are of clinical concern (5, 7). Therefore, it is crucial to predict the stroke outcomes at an early stage (4, 8). In clinical studies, imaging biomarkers such as midline shift (MLS) and swelling volume (4, 8, 9) have been critical predictors of stroke outcomes. They are also crucial criteria in various scoring systems such as EDEMA (10) and TURN scores (11). However, these critical markers can only become visible on brain images days after stroke onset, which delays clinical decisions. Thus far, predicting these markers remains challenging as brain swelling is a complex interaction between swollen tissue and brain bone structures, and brain and lesion anatomy can play a key role in this process (6, 12, 13). To date, (14) has been the only existing study that predicts brain tissue displacement within a single brain geometry under impact, and there is still a lack of research on the prediction of post-stroke brain swelling.

To explore novel tools to predict brain swelling, we propose a computational pipeline where FE models are utilised to generate high-resolution tissue deformation by solving biomechanical governing equations. Using the model, varied 3D brain and lesion geometries can be generated and simulated to obtain 3000 in silico stroke cases. The results are then used to train and test the designed DNN to predict brain swelling with high accuracy. Furthermore, the model is used to process brain images of 60 patients and predict MLS and brain volume change. The predicted imaging markers show a reasonable prediction of disability and death 90 days after stroke. In this study, the first computational workflow for brain swelling prediction after stroke is thus proposed to assist early clinical decision-making.

## Results

### Model Overview

The human brain is modelled using a fivecompartment (arteriole blood, capillary blood, venule blood, interstitial fluid, and tissue displacement) poroelastic theory, which characterises tissue deformation and fluid transfer in the brain vasculature and interstitial space. In lesions with damaged BBB, blood components leak into the interstitial space, and the leakage rate depends on fluid and osmotic pressure gradients. As the key component that determines brain swelling, fluid leakage is represented by incorporating Donnan’s effect (15) into the poroelastic model. This gives us a comprehensive mathematical description of blood flow, interstitial fluid flow, BBB leakage, and tissue deformation during brain swelling (Fig. 1b).

**Fig. 1.**
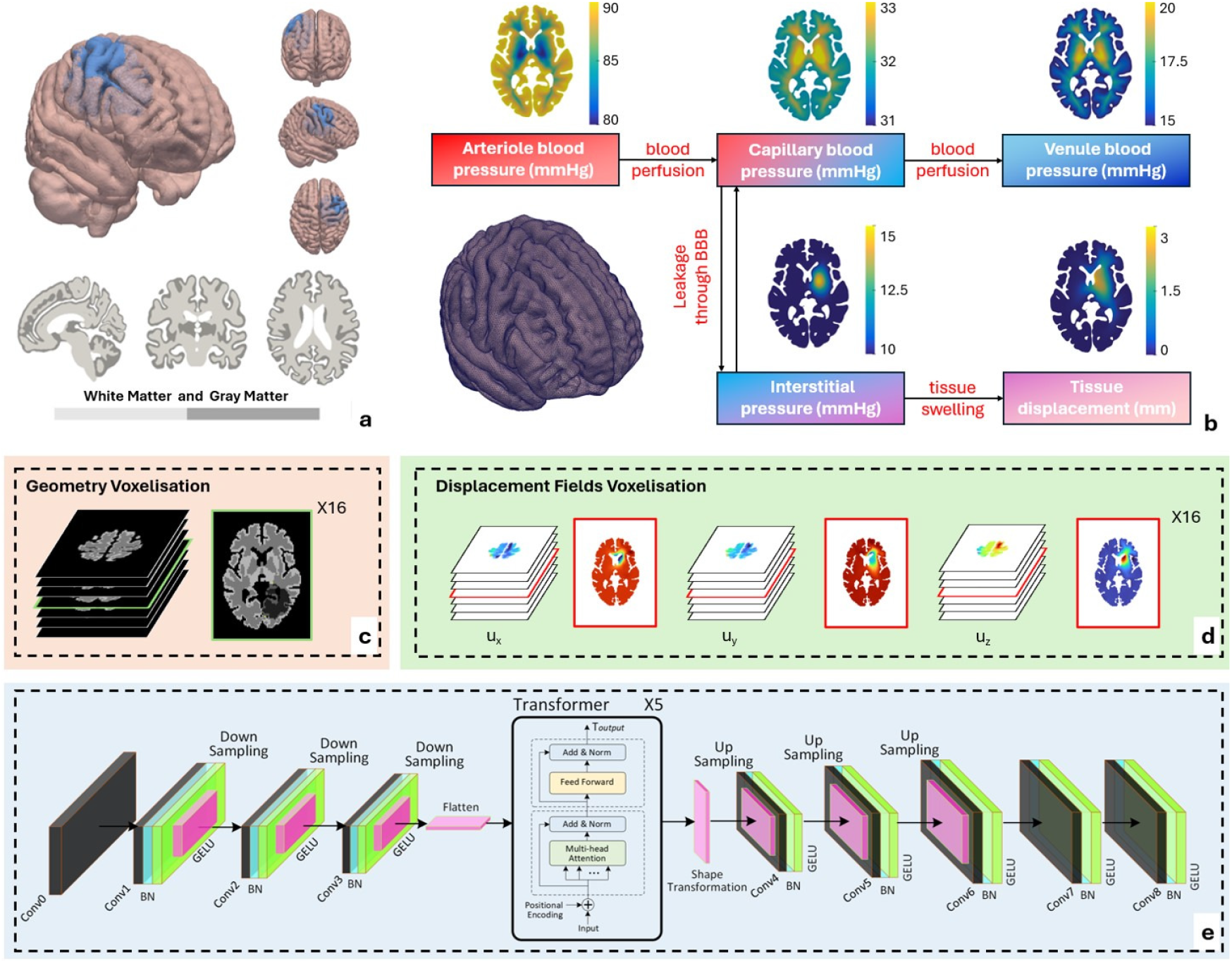
The workflow of the models. (a) The geometry of the brain, lesion and the white and grey matter. This gives us four computational domains. (b) 3D brain mesh. Brain deformation is simulated in the five-compartment poroelastic model. (c) The brain is voxelised and the stroke brain anatomy is used as the input of the DNN. (d) The simulated brain displacement is voxelised and used as the output of the DNN. (e) The input with dimensions of 80 *×* 104 *×* 88 is first processed with Conv0 layer to obtain 16 input channels. In Conv1, 2, 3 & 4 layers, the number of channels is increased by twice, followed by the batch normalisation layers and the GELU activation functions. The downsampling layers halve the three dimensions of the inputs. After down-sampling, the data has 128 channels and is flattened and used for the input of the transformer encoder. Thereafter, the output of the transformer is reshaped to 10 *×* 13 *×* 11 with 128 channels. The DNN then up-samples the data with CNN layers. Conv4, 5, 6 & 7 halve the number of channels whilst the upsampling layers doubles the size of images. Finally, the Conv8 reduces the number of channels to 3 to obtain the displacement fields in three directions. *L*_*p*_ loss has been used for the training of the DNN. The model is trained 800 epochs with a learning rate of 5 *×* 10^*‒*5^ and a decay rate of 0.1. A weight of 20 is imposed on the lesion core region and a weight of 10 is imposed on the right side of the brain. Meanwhile, the background is imposed zero loss (Fig.S8). A background mask is used to remove the final predicted displacements of background voxels.

Brain deformation is simulated using 3D brain meshes with 1.4 million tetrahedral elements. To investigate varied stroke brain anatomy, 30 brain meshes are generated from a population-averaged brain mesh, where affine transformation is utilised to register the brain mesh with the imaging of 30 brains from a clinical cohort (16). The brain mesh is divided into grey and white matter subdomains and different flow and mechanical parameters are applied according to physiological data (12, 17). Thereafter, 100 leaky lesion cores are generated with random locations and sizes in the right hemisphere of each brain. This results in 3000 in silico stroke models, with each model containing four subdomains: healthy white matter, healthy grey matter, lesion white matter, and lesion grey matter (Fig. 1a).

After obtaining simulation results, the continuous FE data are voxelised with a 2 *×* 2 *×* 2mm^3^ sampling to align with the resolution of MRI/CT images (Fig. 1c&d). The training data are 2400 cases in 24 brains, whilst the other 600 cases in 6 brains with varied brain geometries are used for DNN testing. The designed DNN predicts the spatial distribution of tissue displacement in the x, y, and z directions by learning stroke brain anatomy as model input (Fig. 1e). Thereafter, a post-processing algorithm is employed to calculate the swollen brain geometries by using the displacement fields. The swollen brain geometries provide visualisation and comparisons to obtain swelling brain volumes after stroke onset.

### Displacement Prediction

As the displacement of the brain tissue can depend on the locations and the sizes of the lesion cores, six stroke cases are selected from the 600 testing cases to present the predicted brain tissue displacement fields. Here, six cases with large deformation are chosen from six different brains (quantitative evaluation of the geometrical differences are summarised in Table 1), respectively, as this helps illustrate the displacement fields with better clarity (Fig. 2). It is noted that the maximum errors in prediction are well below 1 mm, with most of the errors not exceeding 0.5 mm across the computational domain. Notably, both the patterns for large (Pred *u*_*x*_ over 9 mm in the fourth brain in column 4, row 4) and small deformation (Pred *u*_*z*_ below 1 mm in the first brain in column 10, row 1) in the complex brain anatomy are captured by the DNN accurately.

**Table 1.** Error quantification in the 600 cases across the six brain geometries in the test dataset, with errors presented as *mean ± SD*.

| Brain IDs | Sagittal Lengths (mm) | Coronal Lengths (mm) | Axial Lengths (mm) | Tissue Volumes (ml) | $u_x$ Error (mm) | $u_y$ Error (mm) | $u_z$ Error (mm) |
| --- | --- | --- | --- | --- | --- | --- | --- |
| 24 | 126 | 172 | 160 | 1266.6 | $0.0373 \pm 0.008$ | $0.0352 \pm 0.009$ | $0.0552 \pm 0.005$ |
| 25 | 132 | 168 | 152 | 1253.0 | $0.0373 \pm 0.008$ | $0.0345 \pm 0.010$ | $0.0543 \pm 0.005$ |
| 26 | 122 | 168 | 148 | 1126.2 | $0.0374 \pm 0.009$ | $0.0360 \pm 0.011$ | $0.0539 \pm 0.006$ |
| 27 | 123 | 176 | 156 | 1243.3 | $0.0405 \pm 0.008$ | $0.0378 \pm 0.010$ | $0.0566 \pm 0.005$ |
| 28 | 124 | 168 | 144 | 1106.1 | $0.0370 \pm 0.010$ | $0.0358 \pm 0.010$ | $0.0532 \pm 0.005$ |
| 29 | 126 | 176 | 158 | 1267.6 | $0.0427 \pm 0.010$ | $0.0387 \pm 0.010$ | $0.0573 \pm 0.005$ |

**Fig. 2.**
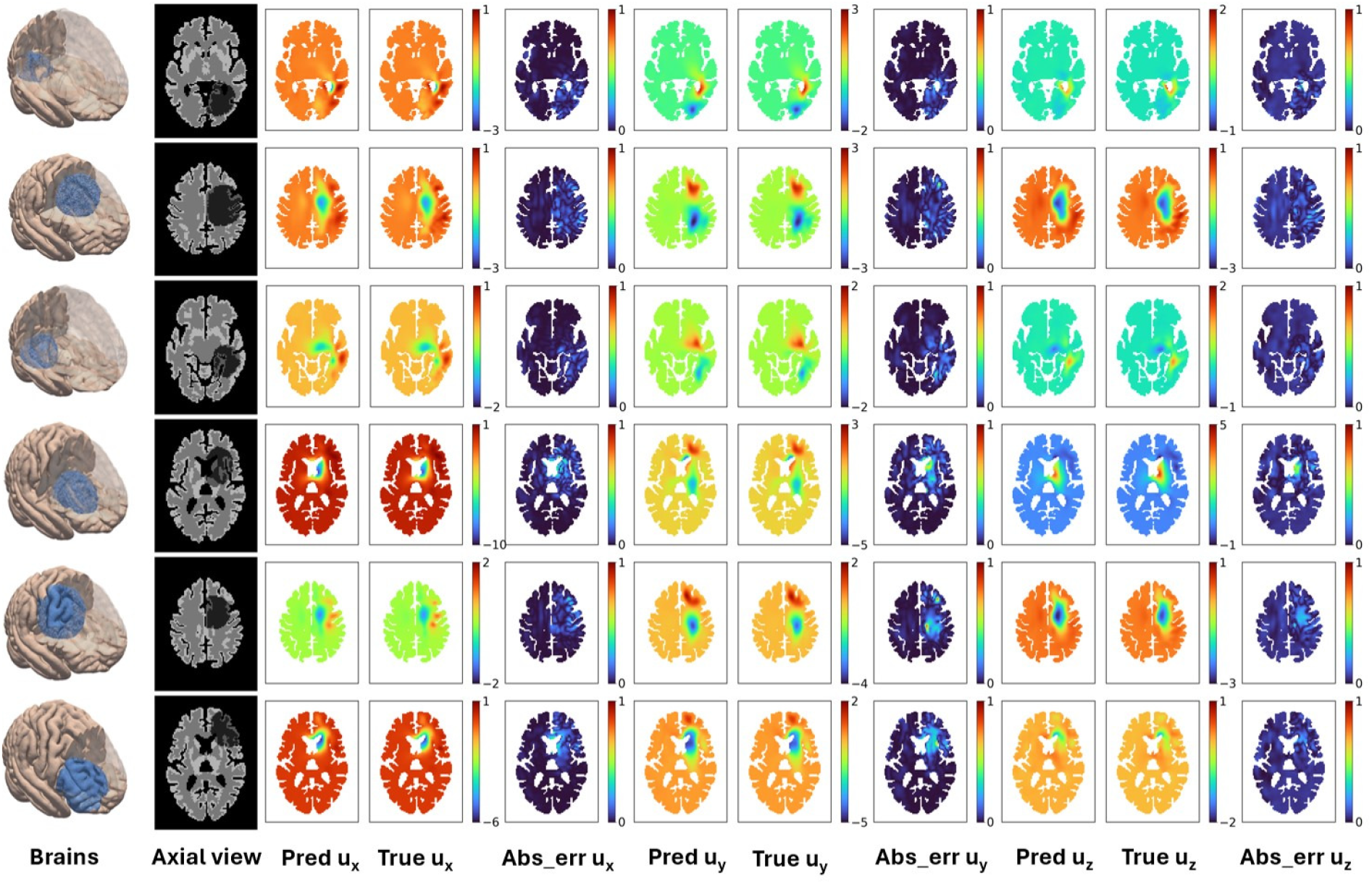
The comparison of the DNN prediction results with the ground truth is illustrated, with all color bars represented in millimeters (mm). The first column is the snapshots of the brain and lesion geometry from the same angle, with the brain segmented to show the stroke locations. Column 2 is the slice of the axial plane across the lesion centroid location. The DNN predicted *u*_*x*_, *u*_*y*_, *u*_*z*_ are shown in columns (3, 6, 9). The FE generated *u*_*x*_, *u*_*y*_, *u*_*z*_ are shown in columns (4, 7, 10), and the errors in prediction are shown in columns (5, 8, 11), respectively. The displacement fields predicted by the DNN are found to achieve nice matches with the FE-generated ground truth. For example, the displacement in x direction majorly concentrates at the front of the ventricle surface in MCA, ACA region lesion (rows 4&6) whilst the displacement is the largest on the back of the ventricle in PCA region lesion (rows 1&3). Lesions at different locations can lead to different patterns of displacement fields, and these features are well captured by the DNN.

### Geometry Prediction

The DNN can predict displacement accurately and the calculated swollen brain geometries show only slight differences when compared with the FE ground truth (Fig.S2-S7). Most prediction errors in tissue-background boundaries are minimal and appear as individual islands (marked in red and green, where red is excessive and green is deficient). This indicates that the errors do not exceed 2 mm and they are acceptable errors that can be caused by the 2 *×* 2 *×* 2mm^3^ voxelisation of brain geometries. The accurate lesion-healthy tissue boundary prediction shows that the lesion swollen volume can be captured by the DNN.

Furthermore, the white-grey matter boundary shows minimal prediction errors, which indicates that the displacement of white and grey matter is well captured. It is worth noting that different mechanical properties are applied to the white and grey matter, where the shear modulus value of white matter is twice of grey matter. This demonstrates that the DNN can learn tissue mechanical properties from simple grey-valued imaging data.

### Error Quantification

To present the prediction outcome of the DNN, brain geometrical features and prediction errors of displacement are summarised in Table 1. The errors in 600 test cases are categorised into six brain geometries, with each group containing 100 cases. Although the brain geometrical features vary (sagittal length, 122-132 mm; coronal length, 168-176 mm; axial length, 144-160 mm; brain volume, 1106.1-1267.6 ml) in the testing cases, the prediction errors are not significantly different between the 600 cases in the six brains. Considering tissue displacements are governed by biophysical laws and are not simple affine transformations of the baseline displacements, these results demonstrate that the geometrical features of the brain have been well learned and extrapolated by the designed DNN.

The RMSE of displacement is found to be much lower than 0.1 mm. Meanwhile, the maximum errors in DNN-predicted displacement are well below 1 mm whilst the spatial resolution of the stroke brain geometry input has a spatial resolution of 2 mm (Fig. 2). This shows that the designed DNN is capable of learning tissue displacement with high accuracy at a subvoxel level using brain imaging data. Importantly, our transformer-based DNN outperforms previous models on FE-based tissue displacement prediction, where the RMSE of displacement prediction is of the order of magnitude of 1 mm (14, 18). Transformer has been the state-of-the-art model in many fields, such as natural language processing and image segmentation (19, 20). These results show that the transformer can also be a powerful tool for learning complex structural features and predicting brain deformation.

### Stroke Outcome Prediction

Thus far, we have obtained a DNN model for the prediction of brain swelling after stroke by using stroke brain anatomy. In this section, we test the capability of the model in generating MLS and brain swelling volumes for an effective prediction of stroke outcomes. Here, we use 60 stroke brain images from the ISLES 2024 open source dataset (21, 22), in which stroke patients brain images, masks of infarct and the modified Rankin Scale (mRS) after 90 days are provided. Of the 60 patients, 31 had poor functional outcomes (mRS*>*2) and 9 died (mRS=6) after stroke. Due to the low-resolution boundaries between the white and grey matter on the brain images, it is challenging to extract the exact boundaries of four subdomains. For simplicity, the centroids and volumes of infarcts are extracted from the dataset to generate lesions of similar locations and sizes in the population-averaged brain (Fig. 3) (16). Thereafter, the stroke brain geometries are used by the DNN to infer MLS and brain volume change for the prediction of disability (mRS*>*2) and death (mRS=6), as shown in Fig. 3.

**Fig. 3.**
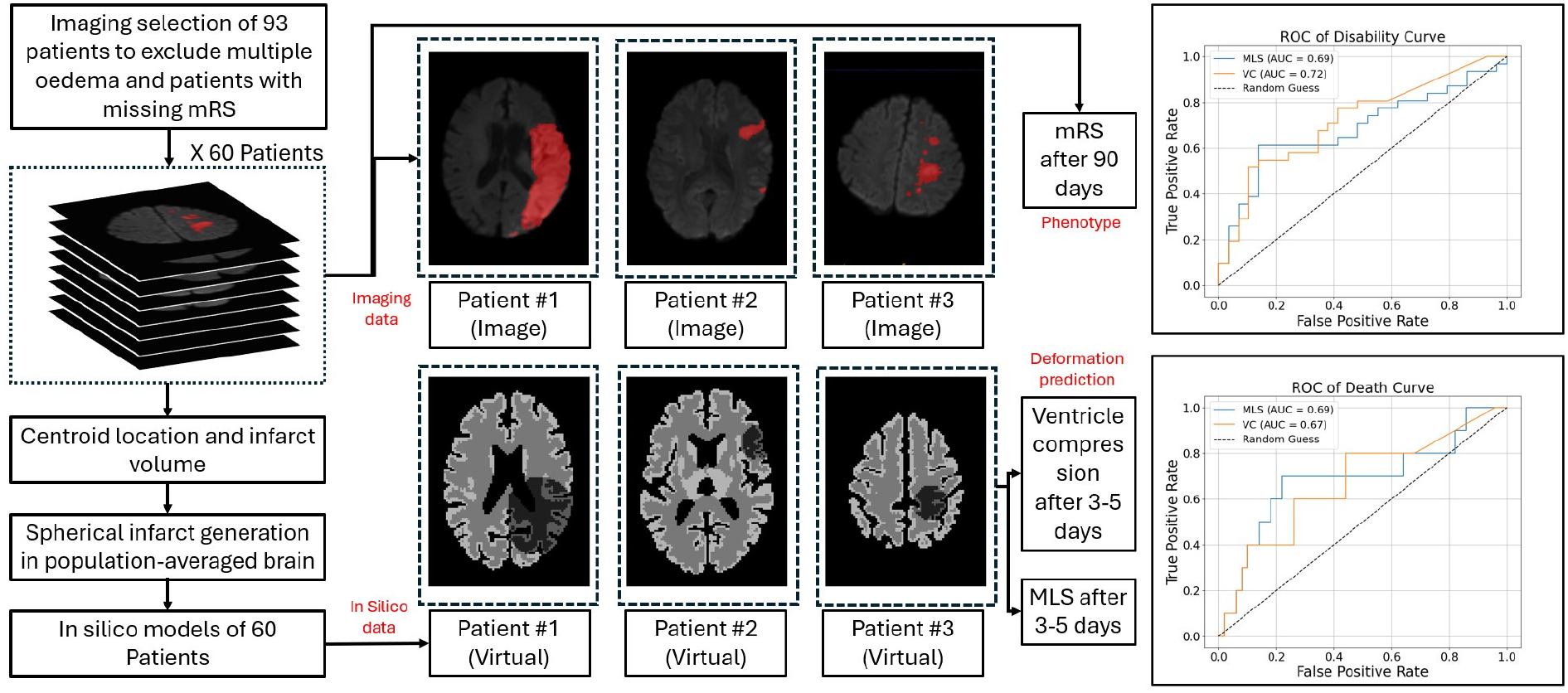
In the ISLES 2024 dataset, 93 patients are initially included, and images with multiple infarcts or missing 90-day mRS are excluded from the analysis. A total of 60 images are selected, and the centroid and volumes of the infarct masks are extracted to create a comparable lesion in the population-average brain model. The computational pipeline is used to obtain midline shift (MLS) and ventricle compression. The resulting predicted radiological markers achieve an AUC of around 0.7, indicating a reasonable prediction of 90-day stroke outcomes.

The predicted MLS (1.38 *±* 1.60 mm) agrees well with previous clinical studies, where MLS is 0.0 (0.0 to 2.0) mm in the good outcome group (mRS *≤* 2) and 2.1 (0.0 to 3.3) mm in poor outcome group after 74.9 (50.5 to 99.6) hours to stroke onset (9). It also agrees with the clinical data (23) where the MLS is 1.6 (0.0 to 2.6) mm 3-5 days after stroke onset. Furthermore, ventricle effacement (2.86 *±* 3.81ml) generated from the DNN agrees well with the data provided in (9), where the change is around 1.61 (−0.62 to 4.69) ml in the good outcome and 3.78 (0.34 to 7.39) ml in poor outcome group. This indicates that the model can predict the key imaging makers that are comparable to real brain deformation after 3-5 days to stroke onset. Furthermore, the AUC is around 0.7 for both MLS and brain swelling volume, demonstrating a relevance between the predicted markers and stroke outcomes. In the results, the AUC of disability is 0.69 for MLS and 0.72 for swelling volume, whereas the AUC of death is 0.69 for MLS and 0.67 for swelling volume. The results match well with the AUC values from (9), where the AUC of disability is 0.68 for MLS and 0.79 for swelling volume and the AUC of death are 0.66 and 0.66, respectively. Furthermore, the predicted MLS and volume thresholds for disability are 1.6 mm and 3.29 ml, which is comparable to the thresholds of 2.0 mm and 3.28 ml in (9). Meanwhile, the thresholds for death are 1.9 mm and 1.34 ml, compared to 3.2 mm and 11.0 ml in (9).

It is noted that the prediction of disability shows a higher AUC than death in the model, this is probably due to that there are only 9 death cases in ISLES 2024. Generally, the prediction shows a nice prediction of stroke outcome with an AUC of around 0.7. This demonstrates that the computational pipeline can be a useful tool in the early prediction of imaging markers and can potentially broaden the time window for clinical decisions after stroke onset.

## Discussions

To the best of our knowledge, this is the first investigation of brain swelling prediction based on learning stroke brain anatomical features. Future application of the DNN tool to predict patient-specific brain swelling will require multi-modal imaging data such as angiography (24) and perfusion MRI (25) for a more comprehensive analysis. Meanwhile, more imaging data of tissue displacement will be needed to understand post-stroke brain swelling. Over decades, significant progress has been made in mapping tissue displacement in the field of brain diseases (26–32). Whilst more patient-specific data are necessary, clinical imaging data are rare and can suffer high noises at the current stage. The computational pipeline demonstrates a novel in silico experimentation to predict brain swelling by learning anatomical features. As tissue deformation in complex anatomical structures can occur in various diseases, such as tumours and haemorrhages, we believe that the proposed approach can inspire the broader field of biomedical studies.

Perceivably, brain swelling is a biomechanical process solely determined by tissue mechanics, lesion swollen volume and boundary conditions. The proposed computational pipeline provides a prediction tool for brain deformation based on our up-to-date mechanical knowledge of the brain. Currently, there is a lack of refined tissue mechanical properties in various brain regions (17), and we thus only use white and grey matter subdomains in the model. The major differentiation in brain tissue mechanical properties has been found between white and grey matter, and experimental evidence shows less difference between other brain regions (17). Therefore, further refinement of brain regions can improve model accuracy but is not expected to significantly affect the prediction outcomes.

There are three major limitations to this study. (1) This study utilises affine transformation (16) to obtain the brain geometries that agree with the clinical brain imaging, and only the centroids and volumes from stroke clinical images are used. This is due to the challenge of obtaining exact subdomain boundaries in low-resolution clinical images. More clinical imaging data can be used to further examine how deformation can vary in different brains. (2) Fixed boundary conditions are imposed on the gyrus, sulcus and falx according to previous mathematical modelling studies (12, 33). Ventricle compression and MLS are usually the signs of malignant stroke, and it is hence reasonable to focus on the ventricle effacement at this stage (8, 12). In clinical settings, however, a swollen brain can also occupy sulci spaces, and a more refined outer boundary condition should be investigated using imaging and clinical data. (3) Although the ventricle compression and MLS agree well with the clinical statistics (Fig. 3), the swollen volumes of the brain tissue can also depend on various biomedical factors, such as collateral score, treatment methods and age (34). Overall, malignant stroke is a complex process, and a comprehensive patient-specific prediction tool will require the joint endeavour of clinicians, bioinformaticians, radiologists and computational biologists.

## Supporting Information Appendix (SI)

Supplemental File includes part of the Results and details of the Methods. The lesion mesh generation, mesh affine transformation, and geometry prediction are in SI Appendix. The loss function is visualised in Fig.S8. Model parameters are shown in Table.S1.

## Materials and Methods

The governing equations of the poroelastic system are given as the following:

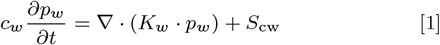

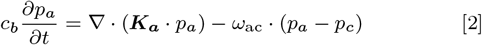

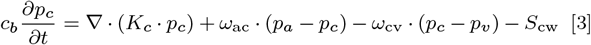

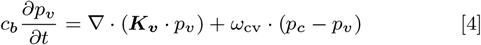

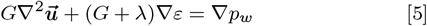

where *c*_*w*_ and *c*_*b*_ are the storage factors of the interstitial fluid and blood in the brain tissue. *a, c, w, v* denote the arteriole, capillary, interstitial space and venule compartments, respectively. The parameters *p*_*i*_ and *K*_*i*_ are the pressure and permeability of *i* compartment. K_***a***_ and K_***v***_ are permeability tensors that align the blood flow direction with penetrating vessels whilst *K*_*w*_ and *K*_*c*_ are isotropic. *ω*_*ij*_ denotes the fluid transfer between *i* and *j* compartments. Meanwhile, *ε* is the dilatational strain and *G* and *λ* are the Lamé constants. The term *S*_cw_ represents the fluid transport from the vasculature into the interstitial space. *S*_cw_ can increase by more than 100 times the normal value (35) in lesion and *S*_cw_ in healthy tissue is thus neglected. The leakage of fluid into the interstitial space *S*_cw_ is derived from modified Starling’s principle and Donnan’s Effect (15, 36):

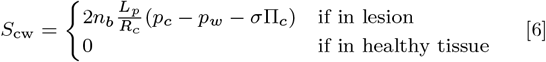

The derivation of *S*_cw_ is shown in (12). *L*_p_ is the hydraulic permeability of the capillary wall, Π_c_ is the osmotic pressure for the plasma components in the blood, *σ* is the reflection coefficient of the original plasma composition, *n*_b_ is the volume fraction of blood vessels in a unit volume of brain tissue, and *R*_c_ denotes the mean vessel radius. The simulations are run 6 time steps with each time step 600 seconds. The final time is shorter than the clinical development time scale as the disruption of BBB is a slow process (12, 37, 38).

Similar to (12), the boundary conditions are: 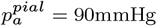 and 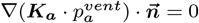 for the arteriole compartment; 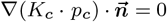 on both the ventricle and pial surfaces for the capillary compartment; 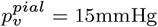 and 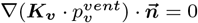 for the venule compartment; 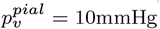 and 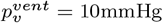 for the interstitial space compartment; 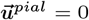 and 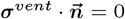 for tissue displacement.

## Supporting information

none

## ACKNOWLEDGMENTS

None.

## References

1. SJ Murphy, DJ Werring, Stroke: causes and clinical features. Medicine 48, 561–566 (2020).

2. PB Gorelick, The global burden of stroke: persistent and disabling. The Lancet Neurol. 18, 417–418 (2019).

3. Y Gu, et al., Cerebral edema after ischemic stroke: Pathophysiology and underlying mechanisms. Front. neuroscience 16, 988283 (2022).

4. S Wu, et al., Early prediction of malignant brain edema after ischemic stroke: a systematic review and meta-analysis. Stroke 49, 2918–2927 (2018).

5. PO Grände, B Romner, Osmotherapy in brain edema: a questionable therapy. J. neurosurgical anesthesiology 24, 407–412 (2012).

6. J Miao, et al., Predictors of malignant cerebral edema in cerebral artery infarction: a meta-analysis. J. neurological sciences 409, 116607 (2020).

7. X Chen, T. Józsa, SJ Payne, Computational modelling of cerebral oedema and osmotherapy following ischaemic stroke. Comput. Biol. Medicine 151, 106226 (2022).

8. ME McKeown, et al., Midline shift greater than 3 mm independently predicts outcome after ischemic stroke. Neurocritical care pp. 1–6 (2022).

9. AC Ostwaldt, et al., Comparative analysis of markers of mass effect after ischemic stroke. J. Neuroimaging 28, 530–534 (2018).

10. CJ Ong, et al., Enhanced detection of edema in malignant anterior circulation stroke (edema) score: a risk prediction tool. Stroke 48, 1969–1972 (2017).

11. D Asuzu, et al., Turn score predicts 24-hour cerebral edema after iv thrombolysis. Neurocritical care 24, 381–388 (2016).

12. X Chen, et al., Modelling midline shift and ventricle collapse in cerebral oedema following acute ischaemic stroke. PLOS Comput. Biol. 20, e1012145 (2024).

13. A Goriely, et al., Mechanics of the brain: perspectives, challenges, and opportunities. Biomech. modeling mechanobiology 14, 931–965 (2015).

14. S Wu, W Zhao, S Ji, Real-time dynamic simulation for highly accurate spatiotemporal brain deformation from impact. Comput. methods applied mechanics engineering 394, 114913 (2022).

15. FG Donnan, The theory of membrane equilibria. Chem. reviews 1, 73–90 (1924).

16. TI Józsa, J Petr, SJ Payne, HJ Mutsaerts, Mri-based parameter inference for cerebral perfusion modelling in health and ischaemic stroke. Comput. Biol. Medicine 166, 107543 (2023).

17. S Budday, TC Ovaert, GA Holzapfel, P Steinmann, E Kuhl, Fifty shades of brain: a review on the mechanical testing and modeling of brain tissue. Arch. Comput. Methods Eng. 27, 1187–1230 (2020).

18. M Pfeiffer, C Riediger, J Weitz, S Speidel, Learning soft tissue behavior of organs for surgical navigation with convolutional neural networks. Int. journal computer assisted radiology surgery 14, 1147–1155 (2019).

19. Y Zhang, H Pei, S Zhen, Q Li, F Liang, Chat generative pre-trained transformer (chatgpt) usage in healthcare. Gastroenterol. & Endosc. 1, 139–143 (2023).

20. K Han, et al., A survey on vision transformer. IEEE transactions on pattern analysis machine intelligence 45, 87–110 (2022).

21. E de la Rosa, et al., Isles’24: Improving final infarct prediction in ischemic stroke using multimodal imaging and clinical data. arXiv preprint arXiv:2408.10966 (2024).

22. EO Riedel, et al., Isles 2024: The first longitudinal multimodal multi-center real-world dataset in (sub-) acute stroke. arXiv preprint arXiv:2408.11142 (2024).

23. HJ Irvine, et al., Reperfusion after ischemic stroke is associated with reduced brain edema. J. Cereb. Blood Flow & Metab. 38, 1807–1817 (2018).

24. R. González, et al., Improved outcome prediction using ct angiography in addition to standard ischemic stroke assessment: results from the stopstroke study. PloS one 7, e30352 (2012).

25. M Grosser, et al., Localized prediction of tissue outcome in acute ischemic stroke patients using diffusion-and perfusion-weighted mri datasets. Plos one 15, e0241917 (2020).

26. M Ferrant, et al., Registration of 3-d intraoperative mr images of the brain using a finite-element biomechanical model. IEEE transactions on medical imaging 20, 1384–1397 (2001).

27. JP Thirion, G Calmon, Deformation analysis to detect and quantify active lesions in three-dimensional medical image sequences. IEEE transactions on medical imaging 18, 429–441 (1999).

28. I Terem, et al., Revealing sub-voxel motions of brain tissue using phase-based amplified mri (amri). Magn. resonance medicine 80, 2549–2559 (2018).

29. C Studholme, C Drapaca, B Iordanova, V Cardenas, Deformation-based mapping of volume change from serial brain mri in the presence of local tissue contrast change. IEEE transactions on Med. Imaging 25, 626–639 (2006).

30. T Hartkens, et al., Measurement and analysis of brain deformation during neurosurgery. IEEE transactions on medical imaging 22, 82–92 (2003).

31. M Soellinger, AK Rutz, S Kozerke, P Boesiger, 3d cine displacement-encoded mri of pulsatile brain motion. Magn. Reson. Medicine: An Off. J. Int. Soc. for Magn. Reson. Medicine 61, 153–162 (2009).

32. D Klatt, CL Johnson, RL Magin, Simultaneous, multidirectional acquisition of displacement fields in magnetic resonance elastography of the in vivo human brain. J. Magn. Reson. Imaging 42, 297–304 (2015).

33. B Tully, Y Ventikos, Cerebral water transport using multiple-network poroelastic theory: application to normal pressure hydrocephalus. J. Fluid Mech. 667, 188–215 (2011).

34. WT Kimberly, et al., Association of reperfusion with brain edema in patients with acute ischemic stroke: a secondary analysis of the mr clean trial. JAMA neurology 75, 453–461 (2018).

35. Y Hakamata, U Ito, S Hanyu, M Yoshida, Long-term high-colloid oncotic therapy for ischemic brain edema in gerbils. Stroke 26, 2149–2153 (1995).

36. BS Elkin, A Ilankovan, B Morrison III, Age-dependent regional mechanical properties of the rat hippocampus and cortex. (2010).

37. GE Lang, D Vella, SL Waters, A Goriely, Mathematical modelling of blood–brain barrier failure and oedema. Math. medicine biology: a journal IMA 34, 391–414 (2017).

38. A Marmarou, et al., Contribution of edema and cerebral blood volume to traumatic brain swelling in head-injured patients. J. neurosurgery 93, 183–193 (2000).

