## Supplementary material for "Early Prediction of Ischaemic Stroke Outcomes: A Novel Computational Approach": none

### PNAS Template for Supporting Information

This PNAS template for Supporting Information (SI) may be used to organize your supporting material. **Once formatted, this first page should be deleted by removing the `\instructionspage` command.** The template is intended to provide a clearly organized PDF file that will ensure readers can easily navigate to sections or specific figures and tables. Movie files or large datasets can be presented as separate files. Further information is available in the PNAS Author Center.

#### Using the template

Specify the title, author list, and corresponding authors with the `\title`, `\author` and `\correspondingauthor` commands. The cover page will be automatically generated with the relevant description of the SI, by the `\maketitle` command.

Figures should be placed on separate pages with legends set immediately below each figure. Table titles should be set immediately above each table. Note that tables extending beyond the width of the page can be included in the PDF or provided as separate dataset files. Oversized/nonstandard page sizes are accepted as part of your SI Appendix file.

References cited in the SI text should be included in a separate reference list at the end of this SI file: (**?** ) and (**?** ).

Supporting information for Brief Reports is limited to extended methods, essential supporting datasets, and videos (no additional tables or figures). Supporting figures and tables are not allowed for Brief Reports.

#### Submitting SI

Delete this first page by removing the `\instructionspage` command, and then save your completed SI file as a PDF for submission. Further submission instructions are available here.

### Supporting Information for

#### Deep Learning Inference of Brain Deformation after Stroke: A Novel Computational Investigation

**Xi Chen, Wahbi El-Bouri, Stephen Payne, Lei Lu**

**Corresponding Author name.**
****

**This PDF file includes:**

Supporting text
Figs. S1 to S8
Table S1
SI References

#### Supporting Information Text

**Lesion Mesh Generation.** The affine transformation method was proposed by (1) and employed to generate brain geometries for oedema simulation in our previous work (2). This generates 30 brain meshes with different sagittal, axial, coronal lengths and volumes. After the meshes are obtained, a random element of the mesh in the right hemisphere is chosen as the starting point of the oedema core and adjacent elements are labelled as the oedema region in a loop of 50-60. As the brain oedema usually does not grow across the hemisphere, the generated oedema core is limited to the right hemisphere of the brain by unlabelling the mesh elements within the left hemisphere. The algorithm for mesh generation in this study is summarised below.

**1.** Generation of brain geometry from population averaged brain mesh using affine transformation

**2.** **a.** Selection of random mesh element in the right hemisphere as the starting point of the oedema core

   **b.** Set iterator $i = 0$ and choose a random loop number between $n = 50 - 60$

   **c.** **IF** $i \leq n$, label adjacent elements to oedema elements as oedema region, $i++$, **ELSE GOTO** Step 3

**3.** Unlabel the mesh elements in the left hemisphere and save the mesh

By using the algorithm, 3000 brain meshes can be obtained in the 30 brains, and 2400 are used for training purposes whilst the rest are employed for the DNN testing. The comparisons of the generated mesh and the brain imaging Fig.S1. Here, we find that using simple affine transformation can avoid the trivial extraction of the brain boundaries and generate brain geometries that have acceptable errors with the patient-specific clinical brain images (detailed comparisons of the 30 brain meshes and clinical images are illustrated in Fig.S1). 2400 oedema cases in 24 brains are used for DNN training whereas 600 cases in 6 brains are used for DNN testing.

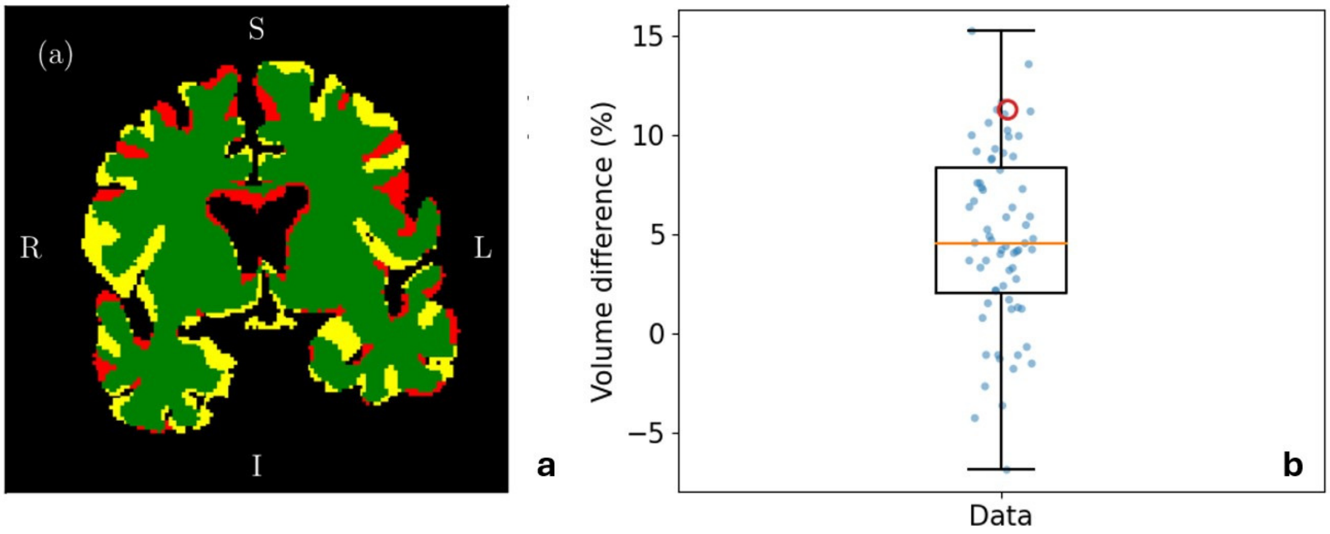


**Fig. S1.** Comparisons of the FE affine transformed brain meshes and the brain volume differences. (a) The comparison of brain geometry with clinical data, reproduced from (1), without change (b) volume difference in percentage, where positive values indicate a larger volume of the FE brains and negative values indicate larger MRI brain volumes. The red marker in (b) is volume difference of the brain geometry shown in (a). It can be seen that the FE generated brain have a reasonably small brain volume difference with clinical brain volume data. Most of the cases have a minimal difference within 10%.

By using the algorithm, 3000 brain meshes can be obtained in the 30 brains, and 2400 are used for training purposes whilst the rest are employed for the DNN testing.

**Model Parameters.** The parameter values and their sources have been presented in our previous studies. In this study, we use the same model parameters as our previous work (2). The only slight change is the mechanical properties of the grey matter and white matter were chosen to be the same in the previous study but were given a difference in shear modulus, where the shear modulus 592.7 for white matter and 296.3 Pa for grey matter. This is important to demonstrate the DNN's capability to capture this difference from the grey values of the FE-generated imaging. The model parameter values are summarised in TableS1.

| Parameter | Value | Parameter | Value |
|---|---|---|---|
| $c_b$ | $1.59 \times 10^{-3}$ Pa$^{-1}$ | $n_b$ | $0.03$ |
| $c_w$ | $3.08 \times 10^{-4}$ Pa$^{-1}$ | $p_v$ | $2000$ Pa $= 15$ mmHg |
| $G_w$ | $592.7$ Pa | $R_c$ | $5 \times 10^{-6}$ m |
| $G_g$ | $296.3$ Pa | $p_a$ | $12000$ Pa $= 90$ mmHg |
| $K_a$ | $1.234$ mm$^3$s kg$^{-1}$ | $\Pi_c$ | $2445$ Pa |
| $K_c$ | $4.28 \times 10^{-3}$ mm$^3$s kg$^{-1}$ | $\omega_{ac}$ | $1.326 \times 10^{-6}$ Pa$^{-1}$s$^{-1}$ |
| $K_v$ | $2.468$ mm$^3$s kg$^{-1}$ | $\omega_{cv}$ | $4.641 \times 10^{-6}$ Pa$^{-1}$s$^{-1}$ |
| $K_w$ | $3.6 \times 10^{-3}$ mm$^3$s kg$^{-1}$ | $\nu$ | $0.35$ |
| $L_p$ | $3.0 \times 10^{-11}$ m/s · Pa | | |

**Table S1. Parameter values**

**Geometry Prediction.** To generate Pytorch tensors for DNN training, the continuous FE computational domains are chosen to be voxelised to generate $80 \times 104 \times 88$ matrices with a spatial resolution of $2 \times 2 \times 2mm^3$, as this is also the resolution of clinical brain MRI images. Similar to the CT/MRI images, grey values of 0, 20, 30, 50 are given to the matrices to represent lesion white, lesion grey, healthy white and healthy grey matter regions. The background of the matrix is given -10. As the tissue deforms, the voxels are moved from their original locations according to the displacement field. In the meantime, the swelling of brain tissue can create void spaces/voxels within the brain tissue regions. We utilise the connectivity of brain tissue voxels to obtain tetrahedral tissue regions and decide if the void voxels are within tissue area after deformation. Finally, the most frequent non-background grey values are used for interpolation to fill the void tissue spaces and create the full post-oedema brain geometries. As the displacement fields are predicted from the brain model with accuracy, the deformation of the brains is thus in good agreement with FE-generated ground truth. The geometrical comparisons are shown in the following figures. Similarly, the brain deformation is marked in pink. The errors in the tissue-background voxels are marked in green and red, whereas errors in lesion-healthy tissue boundaries are marked yellow and blue. Errors in white-grey matter boundaries are marked in orange and purple.

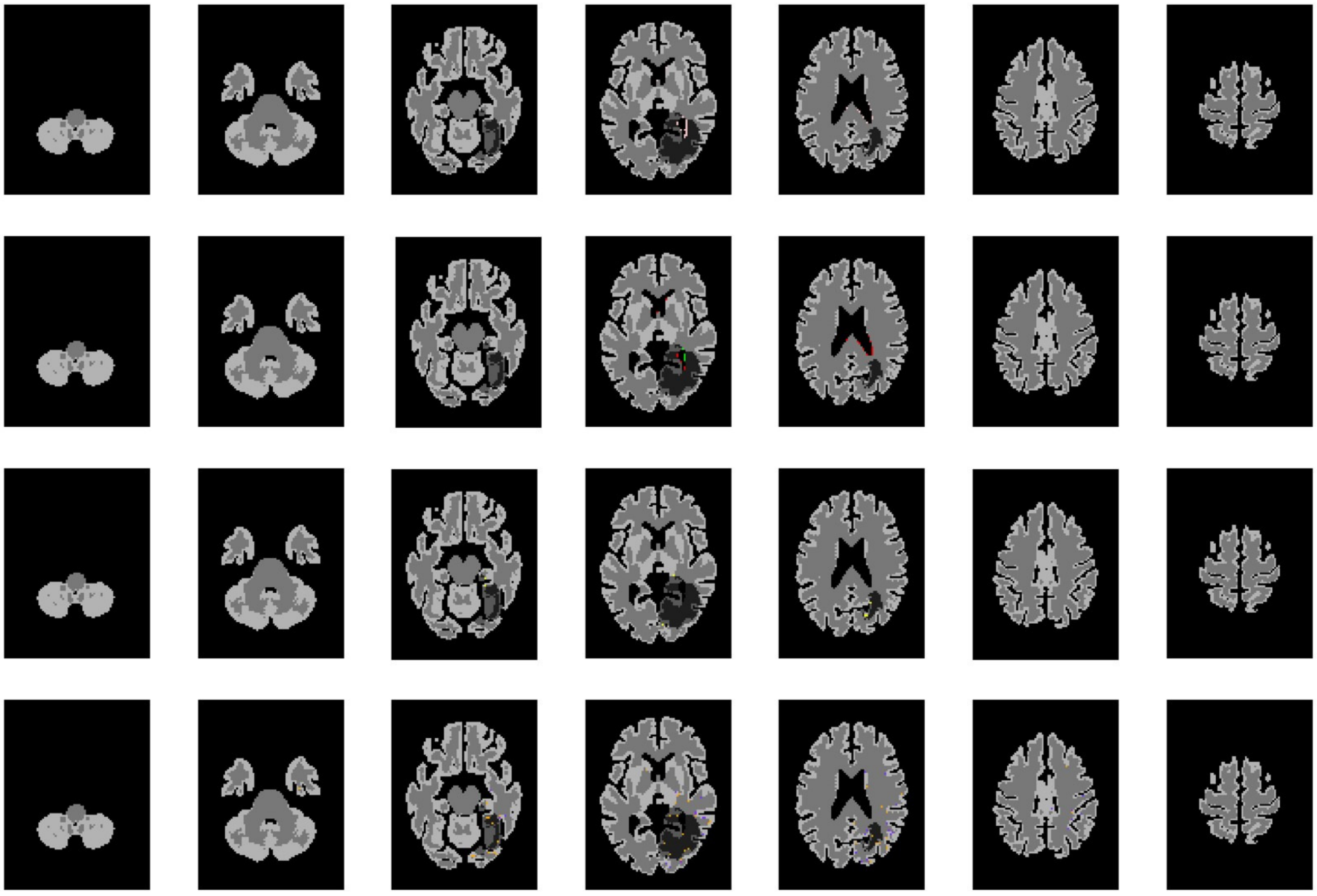

**Fig. S2.** Brain 25, ID 9

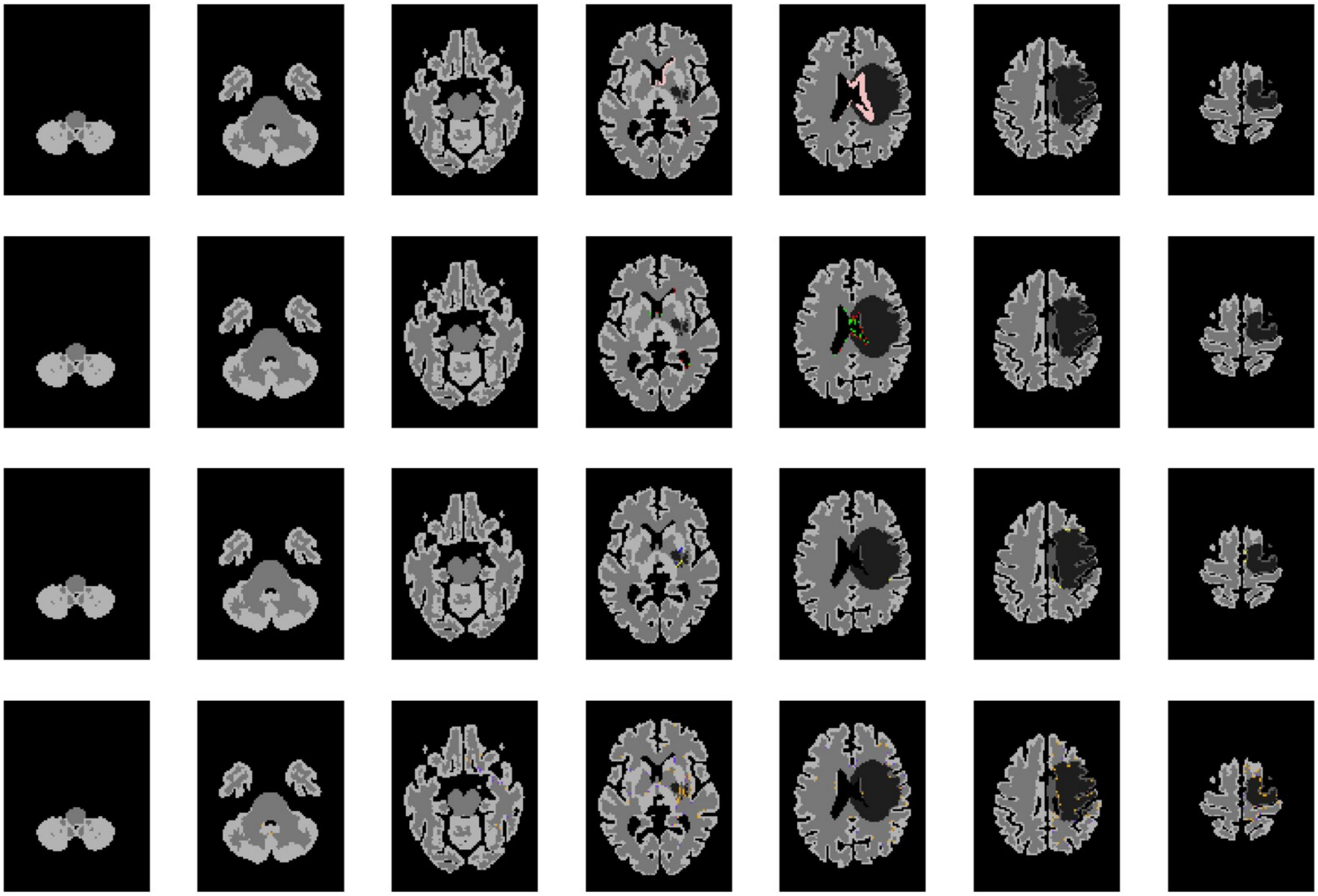

**Fig. S3.** Brain 29, ID 50

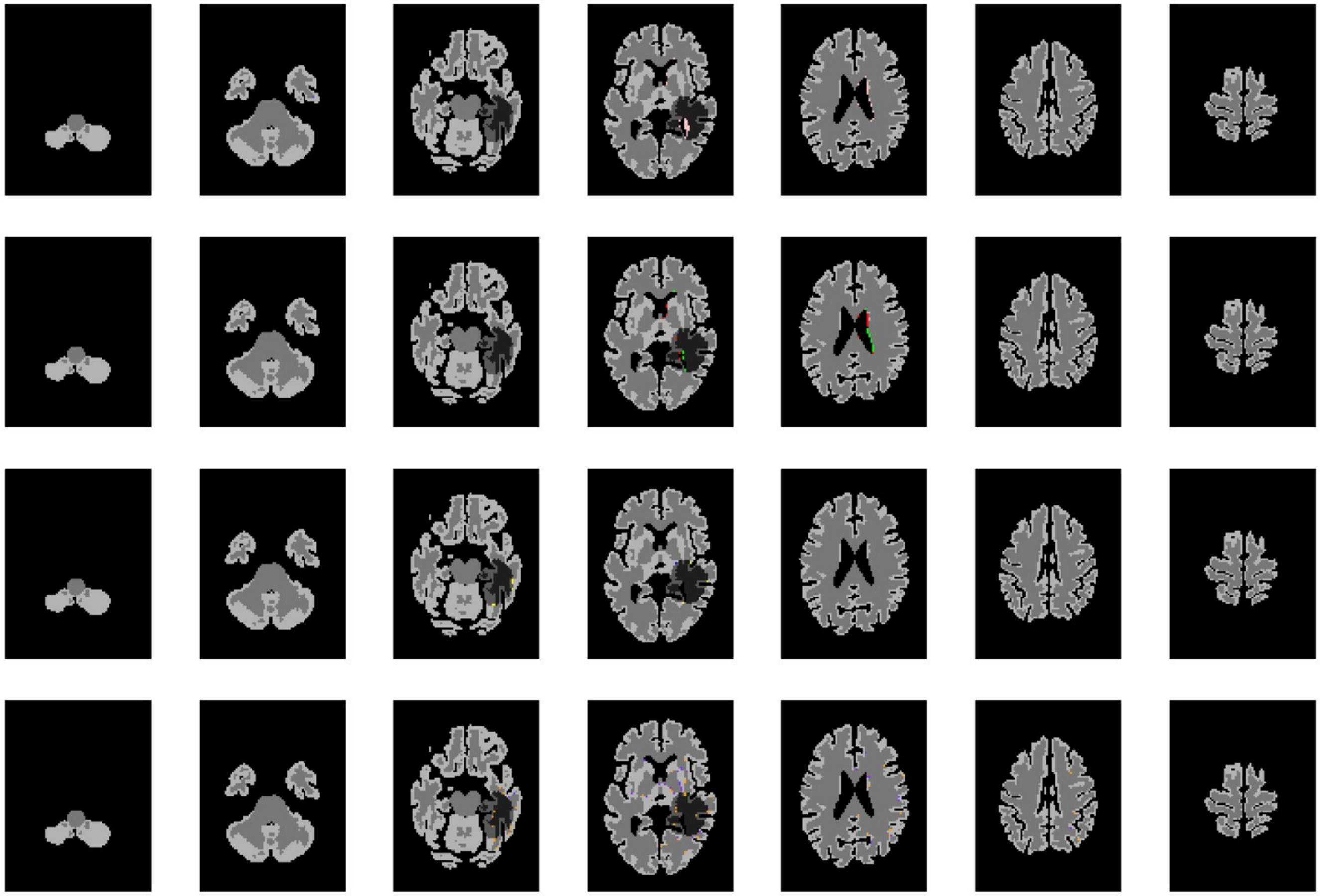

**Fig. S4.** Brain 29, ID 50

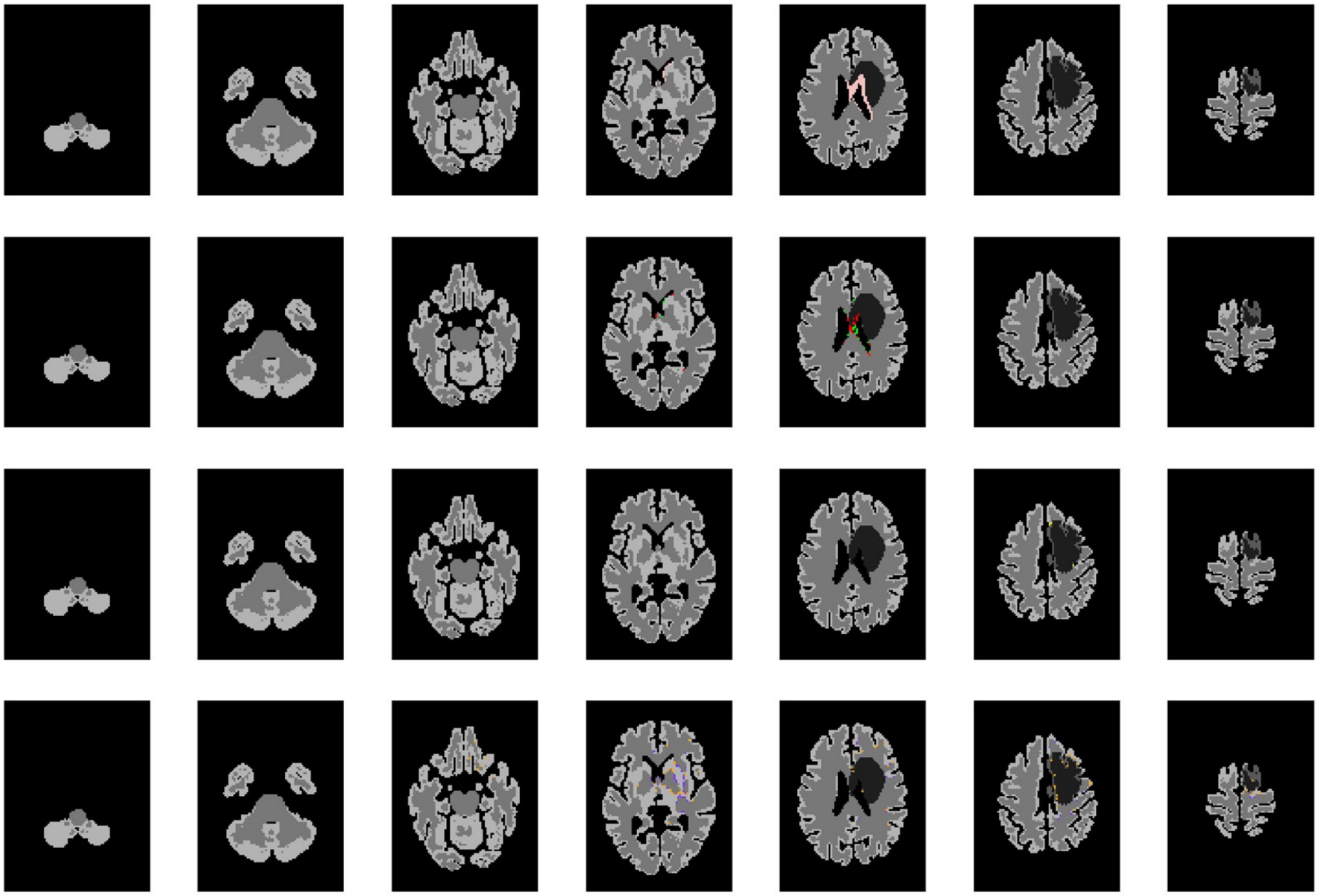

**Fig. S5.** Brain 29, ID 50

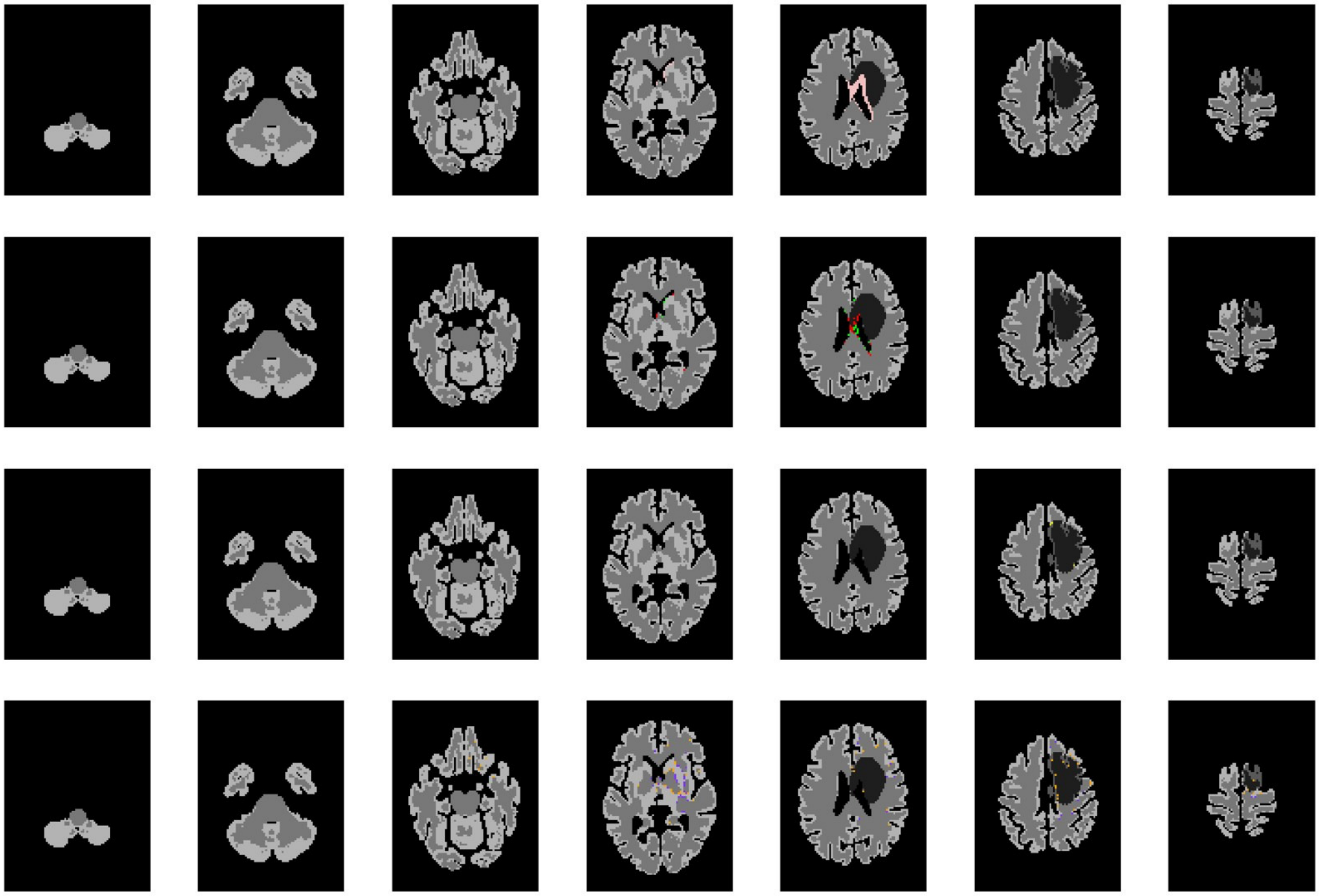

**Fig. S6.** Brain 28, ID 50

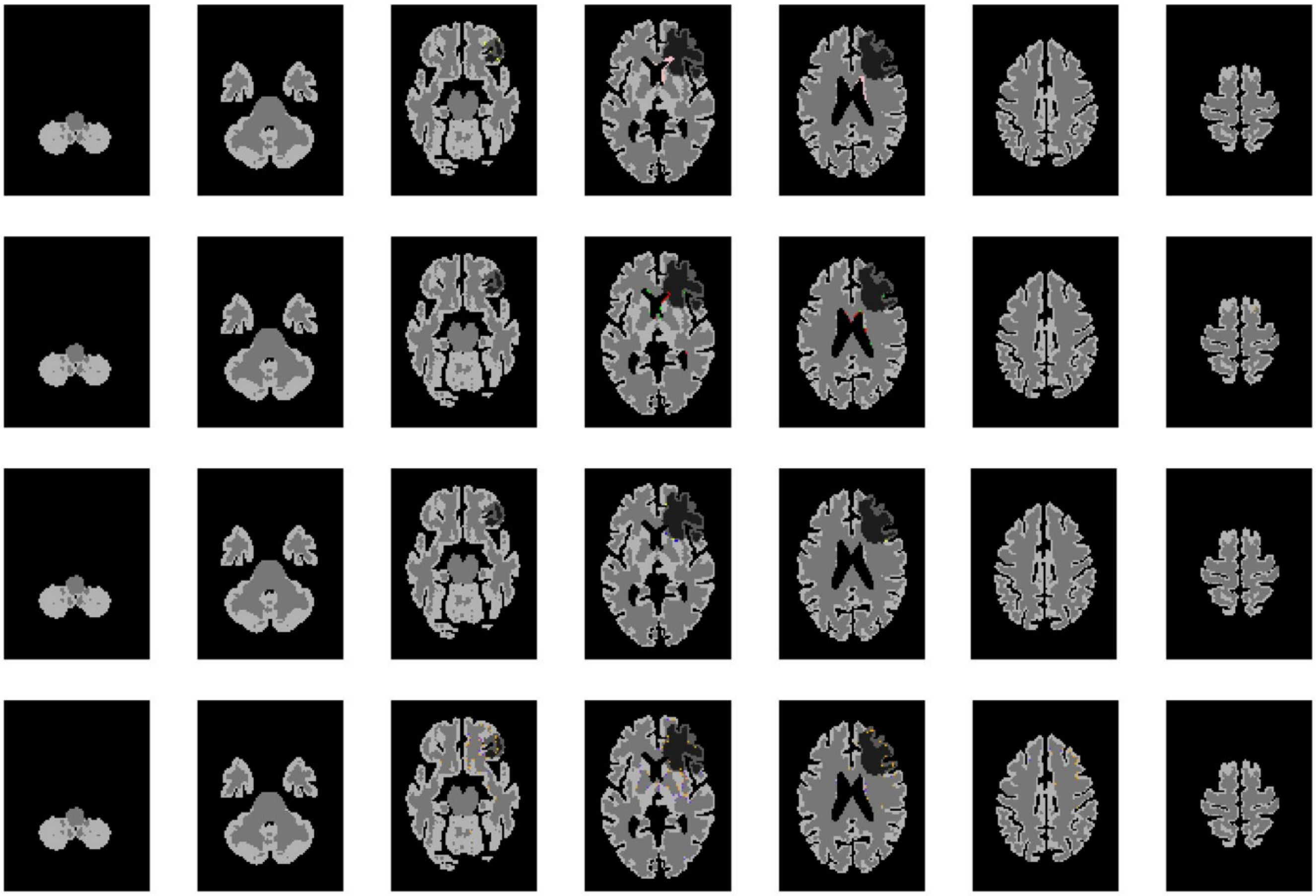

**Fig. S7.** Brain 29, ID 50

**Loss Function Visualisation.** The visualisation of loss weight is presented in Fig.S8.

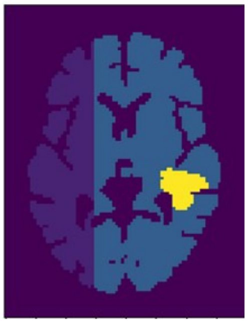

**Fig. S8.** The weight for MSE loss function, where oedema core is weighted by 20, right side of the brain is weighted by 10, and the background loss is imposed zero.

#### References


1. TI Józsa, J Petr, SJ Payne, HJ Mutsaerts, Mri-based parameter inference for cerebral perfusion modelling in health and ischaemic stroke. *Comput. Biol. Medicine* **166**, 107543 (2023).
2. X Chen, et al., Modelling midline shift and ventricle collapse in cerebral oedema following acute ischaemic stroke. *PLOS Comput. Biol.* **20**, e1012145 (2024).
